# Effects of heat-killed *Enterococcus faecalis* T-110 supplementation on gut immunity, gut flora, and intestinal infection in normal aged hamsters

**DOI:** 10.1101/2020.10.05.326124

**Authors:** Takio Inatomi, Konosuke Otomaru

## Abstract

Infectious diseases are a threat to elderly people, whose immune systems become depressed with age. Among the various infectious diseases, *Clostridium difficile* infections in particular lead to significant mortality in elderly humans and are a serious problem worldwide, especially because of the increasing infection rates. Probiotics have been proposed as an effective countermeasure against *C. difficile* infection. The aim of this study was to evaluate the effects of heat-killed *Enterococcus faecalis* T-110 on intestinal immunity, intestinal flora, and intestinal infections, especially *C. difficile* infections, in naturally ageing animals, for extrapolation to elderly human subjects. Twenty female hamsters were randomly distributed into two groups. Group 1 was fed a basal diet, and group 2 was fed a basal diet supplemented with heat-killed *E. faecalis* for 7 days. Heat-killed *E. faecalis* T-110 improved gut immunity and microflora, especially *Clostridium perfringens* and *C. difficile*, of the normal aged hamsters. Heat-killed *E. faecalis* T-110 may, therefore, be a countermeasure against age-related immune dysfunction and intestinal infections, especially *C. difficile* infection, in elderly humans. However, further investigation in humans is needed.

## Introduction

Infectious diseases are a leading cause of mortality and significant morbidity in elderly adult humans, who are at greater risk than younger populations [1]. Increasing age has been associated with diminished humoral and cell-mediated immunity against newly encountered pathogens or vaccines [2-6], which suggests the need for countermeasures against age-related immune dysfunction.

Among the many infectious diseases, *Clostridium difficile* infections are a social problem in elderly humans. These bacteria produce a toxin resulting in symptoms ranging from mild diarrhoea to inflammation of the bowel (pseudomembranous colitis), which can cause death. *Clostridium difficile*-associated diarrhoea is a severe form of diarrhoea caused by *C. difficile* in humans. There are three key risk factors associated with the development of this infection: antibiotic use [7], increasing age [8], and hospitalisation [9]. *Clostridium difficile*-associated diarrhoea is responsible for approximately 10%–20% of all cases of antibiotic-associated diarrhoea [10] and can occur up to 8 weeks after antibiotic therapy [11]. With the increasing threat of *C. difficile* infection, probiotics have been proposed as one of the effective countermeasures against *C. difficile* infection [12-14].

Probiotics have been defined as live, microbial, food components that are beneficial for human health. Recently, because they have been shown to exhibit beneficial effects equal to those of live microbes; genetically engineered microbes and nonviable microbes are regarded as probiotics [15,16]. Lactic acid bacteria, one of the most common types of probiotic bacteria, have been reported to exhibit beneficial effects on host homeostasis, including activation of the immune function [17,18]. To date, many heat-killed lactic acid bacteria have been shown to modulate specific and/or non-specific immune responses in animal models and occasionally in human subjects [19].

Regarding the use of probiotics for infection control, it is important that the probiotics administered are not infectious. *Enterococcus faecalis* T-110 (TOA Pharmaceutical Co. Ltd., Tokyo, Japan), which belongs to the group of lactic acid bacteria, is unlikely to be a causative agent of opportunistic infection [20]. *Enterococcus faecalis* T-110 is approved for medical use and is widely used in Japan, China, and India for the treatment and prevention of infectious diseases. In terms of safety, *E. faecalis* T-110 is considered suitable for treating and preventing infectious diseases.

Several studies have shown that the ageing process affects the gut flora [21-24]. Generally, senescence-accelerated animals are often used to investigate the effects of ageing. However, few studies have shown that the gut flora of senescence-accelerated animals is similar to that of normal, ageing animals. Stephan et al. [25] reported that laboratory mice have a different gut microbiota than free-living mammals and humans, making them unsuitable for the study of gut immunity. *Clostridium difficile* is a bacterium endemic to the intestine of hamsters. Elderly hamsters often suffer from diarrheal infections, especially *C. difficile* infections [26]. These factors suggest that they are the best model of *C. difficile* infections in elderly animals. Challenge tests are generally conducted in bacterial infection tests, but these are considered to be unsuitable as infection models for indigenous intestinal bacteria caused by immunosuppression due to ageing. Few studies have investigated the effects of safety-guaranteed, heat-killed bacteria on intestinal immunity, gut flora, or intestinal infections in normally aged animals. Therefore, the aim of this study was to evaluate the effects of heat-killed *E. faecalis* T-110 on intestinal immunity, flora, and infections in naturally ageing animals, for prospective extrapolation of such information to studies on elderly humans.

## Materials and Methods

### Ethical approval

This study was conducted at Inatomi Animal Clinic in Tokyo Prefecture, Japan. It was performed under the fundamental guidelines for the proper conduct of animal experiments and related activities at academic research institutions under the jurisdiction of the Ministry of Education, Culture, Sports, Science and Technology. It was approved by the Ethics Committee of the Inatomi Animal Clinic (Tokyo, Japan; approval number 2020-001).

### Animals, diets, and management

A total of 20 healthy, 547-day-old female hamsters (*Phodopus sungorus*) were purchased from Japan SLC, Inc., Hamamatsu, Japan, and acclimatised for 10 days prior to use in the experiments. These animals were healthy and did not receive any treatments prior to the study. They were randomly divided into two treatment groups (groups 1 and 2) of 10 hamsters each and housed individually in a cage (27 × 15 × 10 cm) in a 24 h light/dark cycle for 14 days. Temperature was maintained at 26 ± 1 °C, and a basic diet (Rodent Diet CE-2, CLEA JAPAN, Tokyo, Japan) and water were provided *ad libitum*.

Group 2 of hamsters received 0.1 ml heat-killed *E. faecalis* T-110 saline suspensions (1.0 × 10^7^ cfu/ml) daily from days 1 to 7. Heat-killed *E. faecalis* T-110, a commercial product of a heat-killed and dried cell preparation (TOA Biopharma, Tokyo, Japan), was used. The heat-killed *E. faecalis* T-110 saline suspensions were prepared, as previously described [27]. The faeces from individual hamsters was checked daily during this experiment and was categorised according to a faecal score (0, normal faeces; 1, loose stool; 2, moderate diarrhoea; 3, severe diarrhoea).

### Immunological study

On days 1, 7, and 14, the faeces of all hamsters were also measured for total immunoglobulin A (IgA) concentration using a commercial enzyme-linked immunosorbent assay (ELISA) kit (Hamster Immunoglobulin A ELISA Kit, My BioSource, Inc, California, USA). The ELISA procedure was conducted according to the protocol of the manufacturer.

### Microbiological study

The faeces of hamsters were used for the microbiological study. Bacterial genomic DNA from the samples was extracted using a commercial extraction system (QuickGene 810 and Quick Gene DNA tissue kit; KURABO, Osaka, Japan), as previously described [28]. Quantitative real-time polymerase chain reaction (PCR) analyses of *Bifidobacterium* sp., *Clostridium perfringens, Lactobacillus* spp., and *C. difficile* were performed using the Rotor-Gene system 6200 (Qiagen, Tokyo, Japan), as previously described [29]. The primer sequences and PCR conditions for each bacterium are given in Table 1.

**Table 1.**
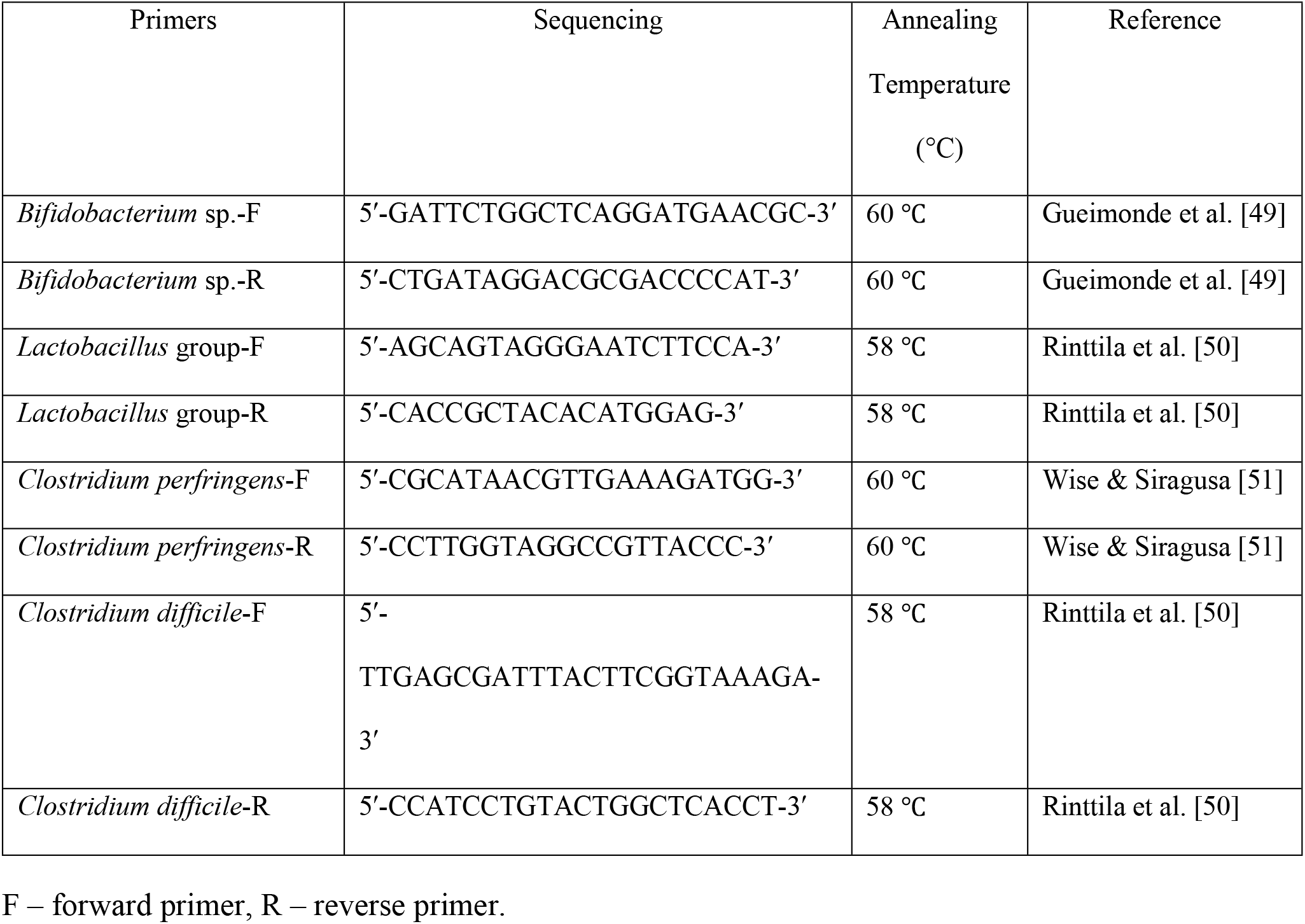
Primers and thermal cycling profiles in this study

### Statistical analysis

Values are presented as means ±standard errors. The Mann–Whitney U-test was applied to analyse differences between mean values in all parameters. Differences between mean values were considered significant at *P* < 0.05 in all statistical analyses. The Mann–Whitney U-test was performed using EZR software (Saitama Medical Center, Jichi Medical University); EZR is a graphical user interface for R (The R Foundation for Statistical Computing, version 2.13.0) [30]. The significance level was set at *P* < 0.05.

## Results

### Total number of days of abnormal defaecation

From days 1 to 7, the total number of days in which abnormal faeces were observed improved in group 2 compared with group 1 (*P* < 0.05) (Table 2). In the second week, there was no difference in the total number of days in which abnormal faeces was detected in group 1 and group 2.

**Table 2.** Total number of days in which abnormal faeces were apparent in hamsters fed a basal diet (group 1) and a 1.0 × 10^**7**^ cfu/ml supplement of heat-killed *Enterococcus faecalis* (group 2)

|  | Days 1–7 | Days 8–14 |
| --- | --- | --- |
| Group 1 | 1.4 ± 0.3<sup>a</sup> | 1.2 ± 0.2 |
| Group 2 | 0.3 ± 0.2<sup>b</sup> | 0.9 ± 0.2 |

<sup>a, b</sup> Different letters within columns indicate differences between treatment groups ( $P < 0.05$ ).

### Immunological study

Total immunoglobulin A (IgA) concentration of faeces was significantly higher in group 2 than that in group 1 at day 7 (Table 3). At days 1 and 14, no difference in total immunoglobulin A (IgA) concentration was detected between group 1 and group 2.

**Table 3.** Total immunoglobulin A (IgA) concentration (mg/g) of faeces in hamsters fed a basal diet (group 1) and a 1.0 × 10^7^ cfu/ml supplement of heat-killed *Enterococcus faecalis* (group 2)

|  | Day 1 | Day 7 | Day 14 |
| --- | --- | --- | --- |
| Group 1 | 2.12 ± 0.09 | 1.98 ± 0.08<sup>a</sup> | 2.01 ± 0.12 |
| Group 2 | 2.08 ± 0.09 | 2.31 ± 0.08<sup>b</sup> | 1.99 ± 0.11 |

<sup>a, b</sup> Different letters within columns indicate differences between treatment groups ( $P < 0.05$ ).

### Microbiological study

On the first day, the numbers of *Bifidobacterium* sp., *C. perfringens, Lactobacillus* spp., and *C. difficile* in faeces were not significantly different between the two groups (Table 4). On day 7, the number of *C. perfringens* and *C. difficile* in faeces was lower in the group 2. The numbers of *Bifidobacterium* sp. and *Lactobacillus* spp. were not significantly different between the two groups. After day 14, the numbers of *Bifidobacterium* sp., *C. perfringens, Lactobacillus* spp., and *C. difficile* was similar in faeces of the two groups.

**Table 4.** Microbiological analyses of faeces (log cells/g) of hamsters fed a basal diet (group 1) and a 1.0 × 10^7^ cfu/ml supplement of heat-killed *Enterococcus faecalis* (group 2)

|  | Day | Group 1 | Group 2 |
| --- | --- | --- | --- |
| *Lactobacillus* spp. | 1 | $5.72 \pm 0.20$ | $5.62 \pm 0.20$ |
| | 7 | $5.40 \pm 0.16$ | $5.45 \pm 0.20$ |
| | 14 | $5.62 \pm 0.24$ | $5.85 \pm 0.23$ |
| *Bifidobacterium* sp. | 1 | $5.35 \pm 0.12$ | $5.53 \pm 0.21$ |
| | 7 | $5.60 \pm 0.14$ | $5.82 \pm 0.22$ |
| | 14 | $5.61 \pm 0.14$ | $5.88 \pm 0.15$ |
| *Clostridium perfringens* | 1 | $5.83 \pm 0.07$ | $5.69 \pm 0.11$ |
| | 7 | $5.72 \pm 0.13^a$ | $4.51 \pm 0.13^b$ |
| | 14 | $5.47 \pm 0.04$ | $5.40 \pm 0.09$ |
| *Clostridium difficile* | 1 | $5.34 \pm 0.05$ | $5.14 \pm 0.10$ |
| | 7 | $5.48 \pm 0.06^a$ | $4.25 \pm 0.09^b$ |
| | 14 | $5.62 \pm 0.06$ | $5.67 \pm 0.06$ |

a, b Different letters within rows indicate differences between treatment groups ( $P < 0.05$ ).

## Discussion

### Immunological study

Immunoglobulin A is one of the main defence elements that prevents pathogenic microorganisms from crossing the intestinal epithelial cell barrier and is important in protecting the intestinal mucosa [31, 32]. In the present study, heat-killed *E. faecalis* T-110 increased total immunoglobulin A (IgA) concentration in the faeces (Table 3). Similar effects have been observed in other studies [33-38]. Havenaar and Spanhaak [39] demonstrated that probiotics stimulate the immunity of animals in two ways: 1) flora from the probiotic migrate throughout the gut wall and multiply to a limited extent and 2) antigens released by dead microorganisms are absorbed and stimulate the immune system. The responsible mechanisms remain unclear; however, it has been suggested that heat-killed *E. faecalis* T-110 stimulates gut immunity, consistent with the results of previous studies.

### Microbiological study

In this study, heat-killed *E. faecalis* T-110 decreased the number of *C. perfringens* and *C. difficile* in aged hamsters (Table 4). Similar effects have been noted in other studies [40-47]. Considering the increased faecal IgA in the current study, it is likely that heat-killed *E. faecalis* T-110 decreased the number of *C. perfringens* and *C. difficile* by improving gut immunity of ageing animals, consistent with the results of previous studies.

### Total number of days of abnormal defaecation

In this study, heat-killed *E. faecalis* T-110 decreased the total number of days of abnormal defaecation. *Clostridium perfringens* and *C. difficile* cause diarrhoea in hamsters [48]. Considering the decreased number of *C. perfringens* and *C. difficile* in the current study, it is likely that heat-killed *E. faecalis* T-110 decreased the total number of days of abnormal defaecation by improving gut immunity in ageing animals.

## Conclusions

Administration of heat-killed *E. faecalis* T-110 improved the gut immunity and flora in normal, ageing animals. However, further elucidation of the mechanism underlying improved immunity is needed.

## Author Contributions

T.I. performed the experiments and drafted the manuscript. K.O. designed the study and supervised the project.

## Data availability

All relevant data are within the paper and its Supporting Information files.

